# The ClpX chaperone and a hypermorphic FtsA variant with impaired self-interaction are mutually compensatory for coordinating *Staphylococcus aureus* cell division

**DOI:** 10.1101/2023.08.22.554348

**Authors:** Camilla Henriksen, Kristoffer T. Bæk, Katarzyna Wacnik, Clement Gallay, Jan-Willem Veening, Simon J. Foster, Dorte Frees

**Affiliations:** Department of Veterinary and Animal Disease, University of Copenhagen, Frederiksberg, Denmark; School of Biosciences, University of Sheffield, Sheffield, United Kingdom; Department of Fundamental Microbiology, University of Lausanne, Lausanne, Switzerland

**Keywords:** *Staphylococcus aureus*, cell division, AAA+ ATPases, ClpX, FtsA, peptidoglycan

## Abstract

Bacterial cell division requires the coordinated assembly and disassembly of a large protein complex called the divisome, however, the exact role of molecular chaperones in this critical process remains unclear. In the important pathogenic bacterium *Staphylococcus aureus*, the ClpX chaperone is essential for growth at 30°C and microscopic analyses suggested that ClpX plays a temperature-dependent role in cell division. We here provide genetic evidence that ClpX unfoldase activity is a determinant for proper coordination of cell division by showing that a spontaneous G325V substitution in the ATP-binding domain of the essential FtsA cell division protein rescues growth and septum synthesis in a *Staphylococcus aureus clpX* mutant. The polymerization state of FtsA is thought to control initiation of bacterial septum synthesis and, while restoring the aberrant FtsA dynamics in *clpX* cells, the FtsA_G325V_ variant displayed reduced ability to interact with itself and other cell division proteins. In wild-type cells, the *ftsA_G325V_* allele shared phenotypes with *E. coli* superfission *ftsA* mutants and accelerated the cell cycle, increased the risk of daughter cell lysis, and conferred sensitivity to heat and antibiotics inhibiting cell wall synthesis. Strikingly, lethality was mitigated by spontaneous mutations that inactivate ClpX. Taken together, our results suggest that ClpX promotes septum synthesis by antagonizing FtsA interactions and illuminates the critical role of a protein unfoldase in coordinating bacterial cell division.

**IMPORTANCE:** Essential biological processes, such as cell division, are performed by multiple proteins working together in dynamic functional complexes. In eukaryotic cells, the disassembly of such molecular machines is often assisted by molecular chaperones capable of unfolding proteins. The ClpX unfoldase is conserved from bacteria to humans, however, the roles of ClpX in bacterial cell biology remain relatively unexplored. By combining genetic methods with super-resolution microscopy techniques, we show here that ClpX and a mutant variant of the essential cell division protein FtsA mutually compensate for each other in controlling cell division of the pathogenic bacterium *Staphylococcus aureus.* The selected FtsA_G325V_ variant has diminished self-interactions and restored the aberrant FtsA dynamics in *clpX* cells suggesting that ClpX promotes cell division by antagonizing FtsA protein interactions. This study, for the first time, illuminates the important role of protein unfoldases in the functioning of the divisome, a multienzyme complex fundamental to bacterial reproduction.

## Introduction

ATP-dependent chaperones are universally required for the folding of nascent polypeptides, and for promoting disassembly of protein complexes and protein aggregates (Saibil, 2013; Mogk *et al.,* 2018). Additionally, a subset of chaperones plays an essential role in protein turnover by targeting protein substrates for proteolytic degradation (van den Boom and Meyer, 2018). One important example is the protein unfoldase ClpX that recognizes and unfolds substrates for degradation by the separately encoded ClpP proteolytic subunit (Olivares *et al.,* 2016). ClpX belongs to the diverse superfamily of AAA+ ATPases involved in various molecular remodeling activities and is highly conserved between eubacteria and mitochondria of eukaryotic cells (Olivares *et al.,* 2016). In mitochondria, the ClpX unfoldase, independently of ClpP, controls key cellular processes such as heme biosynthesis (Kardon *et al.,* 2020) and nucleoid distribution (Kasashima *et al.,* 2012). So far, the ClpP-independent roles of ClpX in bacterial cell biology remain relatively unexplored.

Bacterial cell division by binary fission requires the coordinated assembly of a large protein complex called the divisome. In almost all bacteria, divisome assembly is orchestrated by the tubulin homolog FtsZ that forms a highly dynamic discontinuous ring-like structure at mid-cell composed of FtsZ protofilaments that treadmill around the division plane (Haeusser and Margolin, 2016; Yang *et al.,* 2017; Bisson-Filho *et al.,* 2017). FtsZ filaments are tethered to the membrane by FtsA, an ATP-binding protein belonging to the actin family. FtsA also plays a crucial role in the hierarchical recruitment of a set of highly conserved cell division proteins and, ultimately, transforming the divisome into a septal peptidoglycan (PG) synthesizing machine (Krupka and Margolin, 2018). In *Escherichia coli*, the essential FtsN is the central trigger of septal PG synthesis; however, the importance of FtsA for activation is evident from the isolation of so-called “superfission” mutations in *ftsA* that bypass the requirement for FtsN (Liu *et al.,* 2015; Tsang *et al.,* 2015).

The opportunistic pathogen *Staphylococcus aureus* colonizes the anterior nares of approximately 30% of the human population as a commensal. This colonization is associated with an increased risk of infections, ranging in severity from skin and soft tissue infections to life-threatening bacteremia and toxic shock syndrome (Wertheim *et al.,* 2005). To divide, *S*. *aureus* builds a septal cross-wall, generating two hemispherical daughter cells that separate by ultra-fast popping when the peripheral septal cell wall is resolved in a process involving the Sle1 cell wall amidase and mechanical crack propagation (Monteiro *et al.,* 2015; Zhou *et al.,* 2015, Lund et al., 2018; Thalsø-Madsen *et al.,* 2020). Sle1 is a substrate of the cytoplasmic ClpXP protease, suggesting that the activity of this important cell wall-degrading enzyme is controlled by regulated proteolysis (Feng *et al.,* 2013). In *S. aureus* cells having the *clpX* gene deleted, the lack of Sle1 degradation resulted in cell lysis at 30°C, but not at 37°C, and it was hypothesized that lysis results from the ClpX unfoldase having a temperature dependent role in septum synthesis that in combination with high Sle1 levels result in lysis as depicted in the model in Fig. 1 (Jensen *et al.,* 2019a). We previously showed that lysis can be prevented by the spontaneous acquisition loss-of-function mutations in *ltaS* (encoding lipoteichoic acid synthase) that rescue the growth of *S. aureus clpX* mutants by reducing the amount of Sle1(Bæk *et al.,* 2016). We reasoned that if aberrant septum synthesis is a defining factor in the compromised growth and lysis phenotype of *clpX* cells, then cultivation of *clpX* cells should also be selective for spontaneous suppressor mutations in cell division genes. Indeed, in this study, we describe the characterization of a fast-growing *clpX* suppressor strain that harbors a single nucleotide polymorphism (SNP) in *ftsA,* introducing a G325V substitution in the ATP-binding pocket of FtsA. The role of FtsA in *S. aureus* cell division has not been investigated previously. Interestingly, we show that the FtsA_G325V_ variant shares phenotypes with *E. coli* superfission *ftsA* mutants as it displays diminished self-interaction and accelerated cell cycle. In wild-type cells, expression of FtsA_G325V_ resulted in lysis of daughter cells and was selective for loss-of-function mutations in the *clpX* gene. Collectively, our data suggest show that ClpX and the FtsA_G325V_ variant mutually compensate for each other to control *S. aureus* cell division and that ClpX stabilizes the activated divisome by antagonizing FtsA protein interactions.

**Figure 1.**
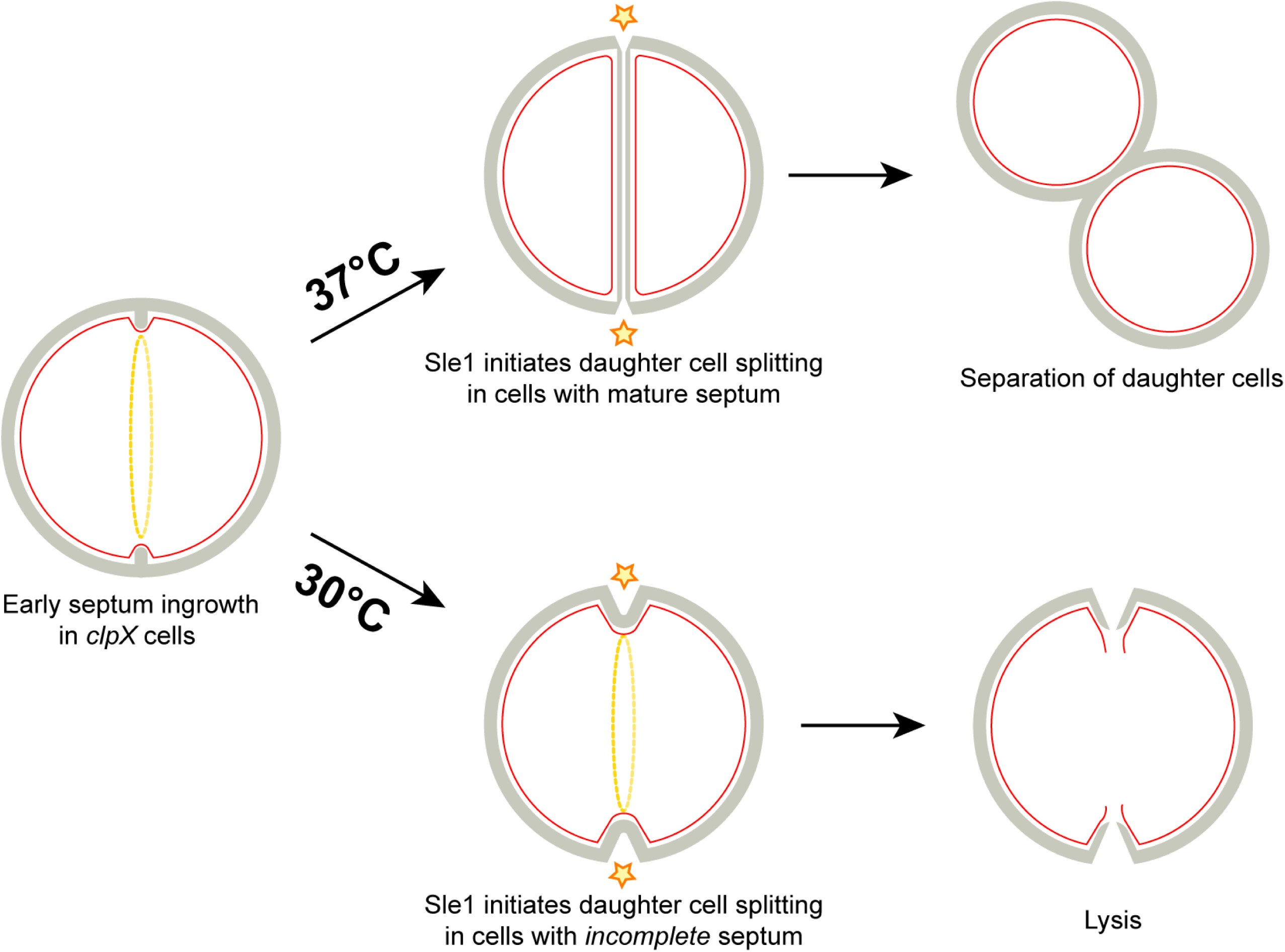
Model of temperature dependent lysis of S*. aureus clpX* mutant. At 37°C, *S. aureus* cells can complete septum synthesis in the absence of ClpX unfoldase activity. Daughter cells separate by ultrafast popping when the Sle1 cell wall amidase cleaves the peripheral septal ring in daughter cells with closed and presplit inner septal cross walls. At 30°C, ClpX becomes critical for inward progression of septum synthesis and the impaired septum synthesis in combination with elevated Sle1 levels result in cells lysis. See text for details. This model is modified from Jensen *et al.,* 2019a.

## Results

### A single amino-acid change in FtsA compensates for the lack of ClpX

When *S. aureus clpX* mutants are incubated at 30°C fast-growing suppressor mutants arise with high frequency and we previously showed that 70% of suppressor mutations inactivated LtaS (Bæk *et al.,* 2016). To identify additional mutations that rescue growth of *S. aureus clpX* mutants, we here picked a *clpX* suppressor mutant (strain 564-30-8) with restored ability to form colonies at 30°C and that retained a wild-type copy of the *ltaS* gene for further characterization (Fig. 2A). The 564-30-8 mutant is a derivative of the clinical strain SA564, a methicillin-sensitive isolate recovered from a patient with toxic shock syndrome (Somerville *et al.,* 2002). Whole-genome sequencing revealed that the 564-30-8 suppressor strain deviated from the parental SA564 *clpX* strain only by a SNP in the *ftsA* gene introducing a G325V substitution in the essential FtsA cell division protein. Western blot analysis revealed that the cellular levels of FtsA are similar between wild-type and *clpX* cells, and that the G325V substitution does not alter the level of FtsA (Fig. 2B). To examine if the *ftsA* mutation was found in other suppressors, the *ftsA* gene was sequenced in 25 *clpX* suppressor strains selected in two different strain backgrounds, SA564clpX and 8325-4clpX (Bæk et al., 2016). One suppressor mutant, strain 83-30-Y selected at 30°C in the 8325-4 background, harbored a mutation in *ftsA* giving rise to the exact same amino-acid substitution (G325V) as the one found in 564-30-8. FtsA is a bacterial homolog of actin, and the mutated G325 and the neighboring G324 are highly conserved among members of the entire actin protein family (Fig. 2C). Importantly, when mapped to the crystal structure of an FtsA dimer, these glycine residues map in a loop structure that forms the hydrophobic binding pocket for ATP, Fig. 2D (Bork *et al.,* 1992, Holmes *et al.,* 1993; Fujita *et al.,* 2014).

**Figure 2.**
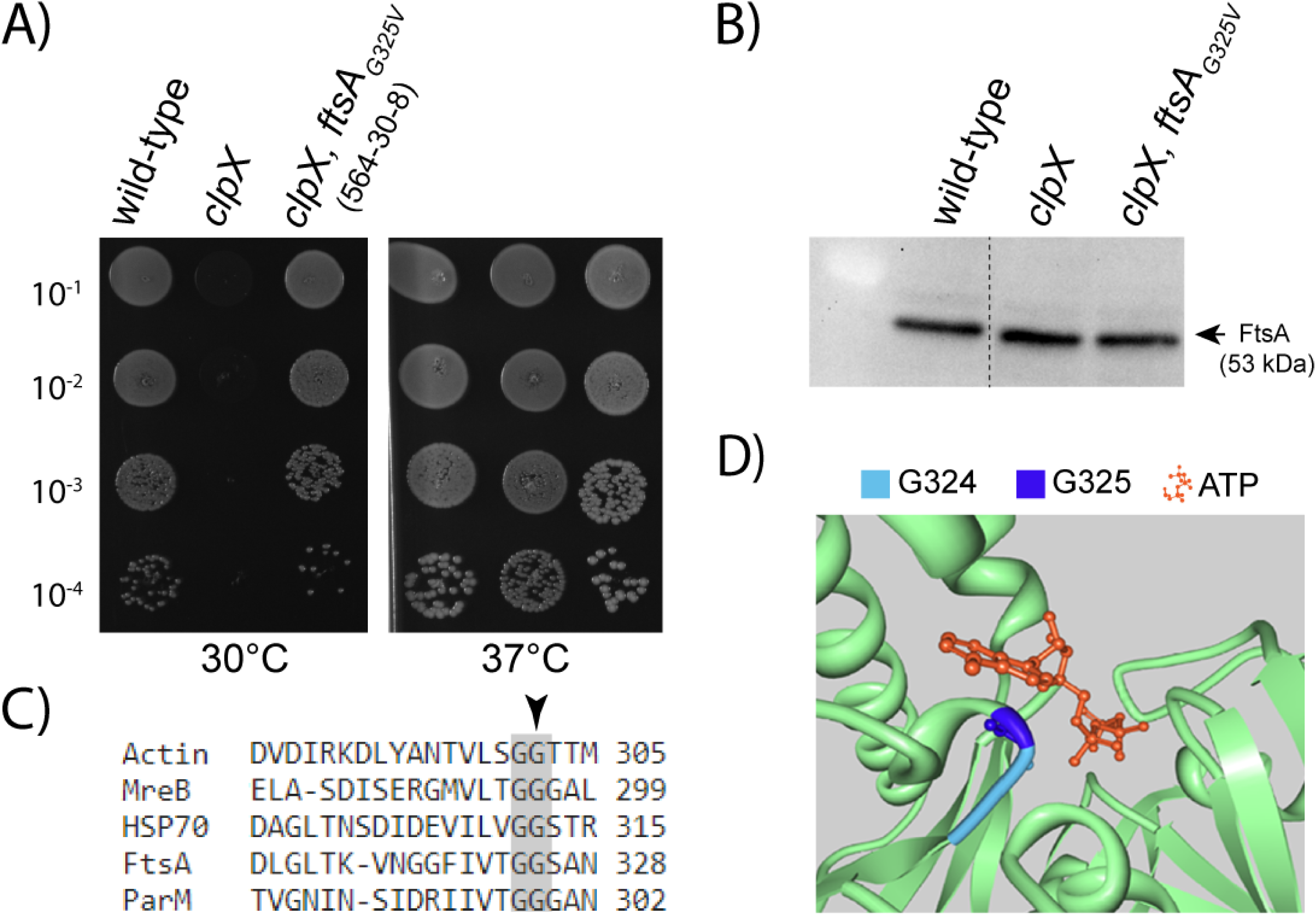
The temperature dependent growth defect of *S. aureus clpX* is rescued by substituting a highly conserved glycine with valine in the ATP binding pocket of FtsA. (**A**) SA564 wild-type strain, SA564clpX, SA564clpX, ltaS, & the suppressor strain 564-30-8 (SA564clpX, ftsA_G325V_) were grown exponentially at 37°C. At OD_600_ = 0.5 ± 0.1, the cultures were diluted 10^1^ to 10^4^-fold and 10 μl of the dilutions were spotted on TSA plates. Plates were incubated at 30°C or 37°C for 24 h. (**B**) FtsA levels in exponential cells of the indicated strains were determined by western blotting using antibodies directed against *Streptococcus pneumoniae* FtsA. Dotted line shows where non-relevant lanes were spliced out of the image. The empty lane to the left contains the protein ladder (**C**) Amino acid sequence alignment of representative members belonging to the actin protein family, (FtsA, HSP70, and ParM from *S. aureus,* MreB from *E. coli*, and human actin). The highly conserved G324 and G325 are highlighted in grey, and the substituted G325 is marked with an arrowhead. (**D**). The mutated glycine is predicted to localize in a loop that forms the hydrophobic binding pocket for the adenosine part of the ATP molecule. Residue G325 is highlighted in dark blue, the neighboring G324 residue is shown in light blue, and a bound ATP molecule in orange in the *S. aureus* FtsA structure (PDB: 3WQT). Visualized with the Protein Workshop (Moreland *et al.,* 2005).

### FtsA_G325V_ prevents spontaneous lysis of *clpX* cells without reducing Sle1 levels

To elucidate how the suppressor mutation rescues the growth of *clpX* cells, we first asked if the *ftsA_G325V_* allele reduces the temperature-dependent lysis of *clpX* cells. The proportion of lysed cells was estimated by staining cultures with propidium iodide (PI), a DNA dye used as an indicator of cell death as it cannot penetrate intact membranes. This analysis revealed that cultures of the *clpX, ftsA_G325V_* suppressor strain contained significantly fewer PI-positive cells than cultures of *clpX* cells when grown at 30°C (3 ± 1.5 % vs 12 ± 1 %, P < 0.001). These results demonstrate that the amino acid change in FtsA counteracts the lysis of *clpX* cells. To examine if the suppressor mutation in *ftsA* prevents lysis of *clpX* cells by reducing Sle1 levels, western blot analysis was performed with protein extracts derived from whole cells and cell walls of wild-type and mutant cells grown exponentially at 30°C or 37°C (Fig. 3A). We additionally examined Sle1 activity using standard zymography (Fig. 3B). For comparison, the previously characterized SA564clpX, *ltaS* suppressor strain was included in the zymographic analysis, and as expected the suppressor mutation in *ltaS* diminished Sle1 activity in *clpX* cells (Fig. 3B; Bæk *et al.,* 2016). In contrast, the elevated Sle1 levels in the SA564ΔclpX cells were not affected by the suppressor mutation in *ftsA* (Fig. 3). We conclude that the *ftsA*_G325V_ allele prevents lysis of *clpX* cells without reducing Sle1 levels.

**Figure 3.**
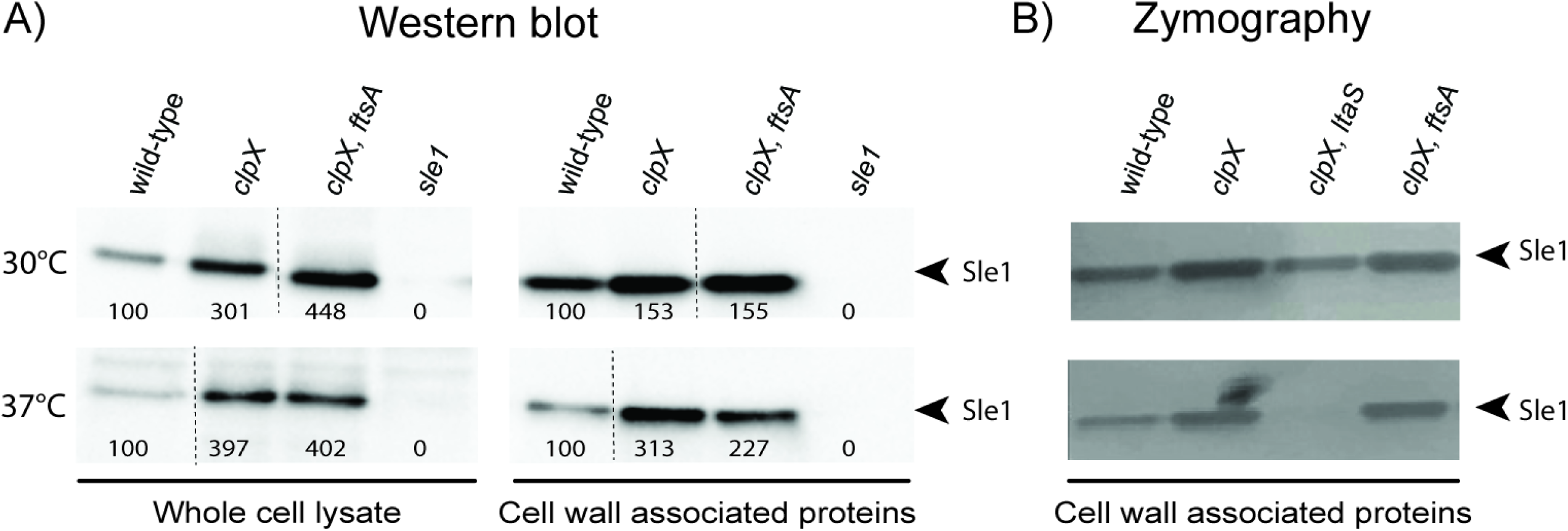
The G325V substitution in FtsA prevents lysis of *S. aureus clpX* mutants without reducing Sle1 levels. (**A**) Sle1 levels in whole-cell lysates and cell wall-associated proteins extracted from SA564 wild-type and the indicated mutants grown exponentially at 30°C or 37°C were determined by Western blotting (representative images of at least two biological replicates). Densitometry measures were acquired using ImageJ and normalized to the wild-type (100%). Dashed black lines show where irrelevant strains were spliced out of the images. (**B**) Zymogram showing autolytic activity from cell wall-associated proteins extracted from SA564 wild-type and the indicated mutants grown exponentially at 30°C or 37°C. Proteins were separated on an SDS gel containing heat-killed *S. aureus* SA564 wild-type cells. The inverted image of one representative gel of two biologically replicates is shown.

### FtsA_G325V_ rescues septum synthesis in *S. aureus clpX* cells

To further investigate how the *ftsA*_G325V_ allele rescues growth of *clpX* cells, we studied the growth of single cells of *S. aureus* SA564 wild-type, SA564clpX, and SA564clpX, ftsA_G325V_ at 30°C using time-lapse phase contrast microscopy. Consistent with published results, the time-lapse experiments demonstrated that only about half of the imaged *clpX* cells (15 of 33 cells) were capable of initiating growth at 30°C (Supplemental Fig. S1), a phenotype that was attributed to ClpX unfoldase activity becoming critical for the progression of septum synthesis as the temperature decreases (Jensen *et al.,* 2019a). In contrast, all imaged *clpX, ftsA_G325V_*cells (45 out of 45) were able to initiate growth at 30°C and increased in numbers almost as fast as wild-type cells, suggesting that the FtsA variant rescues the septum synthesis defect of *clpX* cells (Supplemental Fig. S1).

We next assessed the capability of the cells to complete septum synthesis by scoring cells according to the state of septal ingrowth using SR-SIM images of cells derived from cultures grown at 30°C or 37°C and stained with the membrane dye Nile Red prior to imaging. As observed previously, the fraction of *clpX* cells displaying a closed septum was significantly diminished in cultures grown at 30°C (P < 0.001) but not at 37°C, emphasizing the temperature-dependent role of ClpX in septum synthesis (Fig. 4A). Furthermore, this analysis revealed that expression of the FtsA_G325V_ variant significantly increased the fraction of *clpX* cells displaying a closed septum at 30°C (Fig. 4A). We noted, however, that a large fraction (49 ± 1%) of *clpX, ftsA*_G325V_ cells displayed septal ingrowths progressing asymmetrically from the cell periphery, a phenotype that was also evident in transmission electron microscopy (TEM) images (Fig. 4B). This phenotype was also observed, albeit less frequently, in *S. aureus clpX* cells (20 ± 1%), while the frequency in wild-type cells was very low (< 3%) (Fig. 4B). Strikingly, and in confirmation that the *ftsA* allele does not reduce Sle1 levels, SR-SIM and TEM images revealed that 18 ± 1 % of *clpX, ftsA_G325V_*cells initiated septal splitting from the peripheral wall prior to septum completion (Fig. 4C). This premature splitting phenotype was not observed in wild-type cells but is a hallmark of *clpX* cells (Fig. 4C).

**Figure 4.**
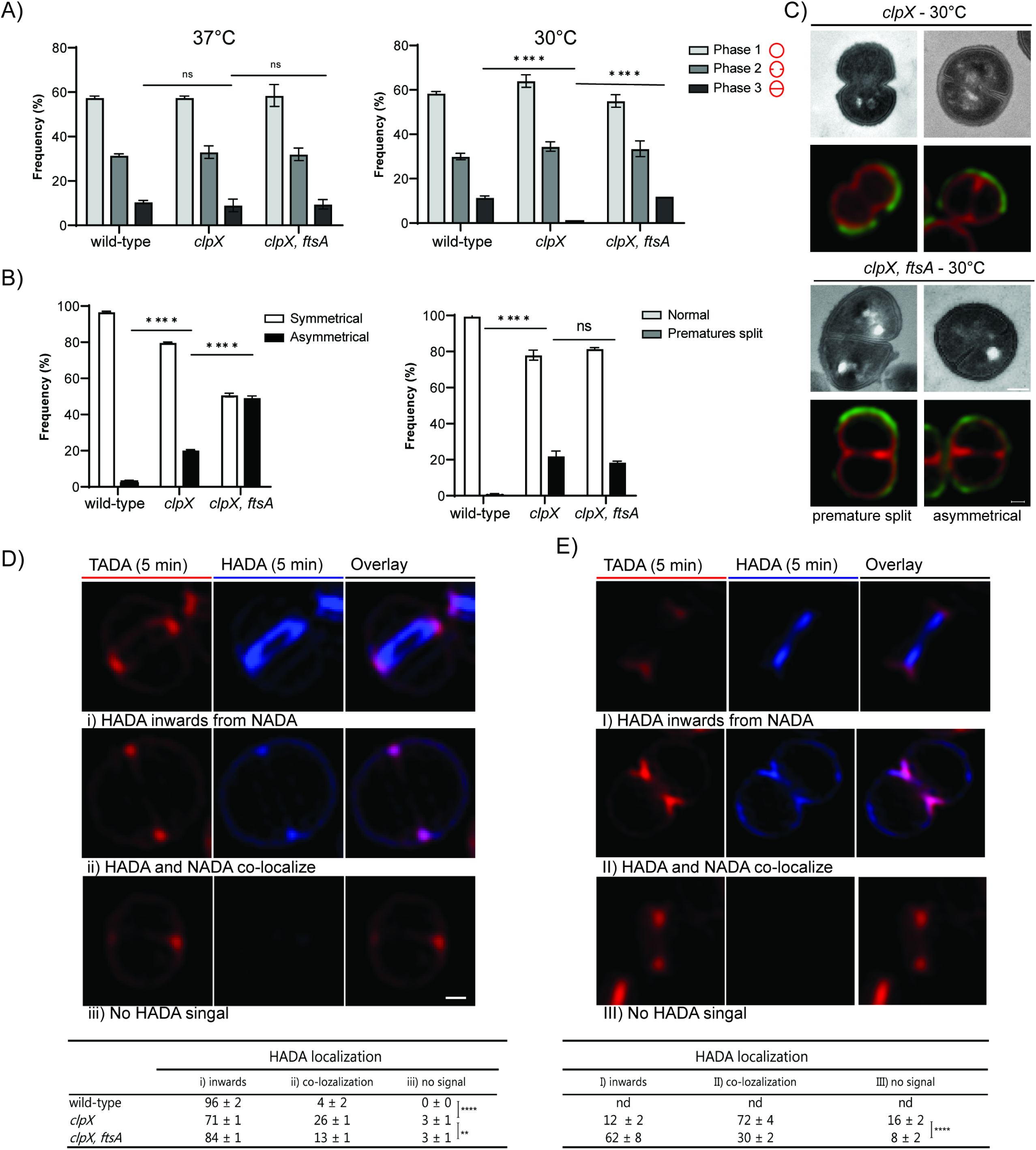
FtsA_G325V_ rescues septum synthesis in *S. aureus clpX* cells. SA564 wild-type, SA564clpX, and SA564clpX, ftsA_G325V_ were grown at 37°C or 30°C as indicated (**A**). 300 cells stained with the membrane dye Nile-Red (red) were scored according to the stage of septum ingrowth: no septum (phase 1), incomplete septum (phase 2), or non-separated cells with complete septum (phase 3) as illustrated; the analysis was based on cells from three biological replicates. For cells grown at 30°C, the fraction of cells displaying asymmetrical ingrowths (**B**) and splitting of daughter cells prior to septum closure (“premature split”, **C**) were scored in 200 septating cells (based on staining with Nile Red; three biological replicates). **** P < 0.0001; ns = non-significant. Statistical analysis was performed using the Chi-square test for independence. (**D+E**) Septal PG synthesis was followed in time by sequentially incubating wild-type, *clpX,* and *clpX; ftsA_G325A_* cells with NADA followed by HADA (10 min labeling each) and determining localization of the HADA signal in cells having NADA incorporated into tiny septal ingrowths being either unsplit (**D**) or showing inward splitting (**E**). Three patterns of NADA/HADA incorporation were observed as illustrated **in D, i-iii** and **E, I-III**, and for each strain the distribution of 50 cells between the three patterns is depicted in the Table below. Frequencies (%) are given as mean and SD for two biological replicates. ** P < 0.01; **** P < 0.0001; ns= non-significant. Statistical analysis was performed using the Chi-square test for independence. Scale bars, 0.2 µm.

To study progression of septal PG synthesis more directly, the three strains (wild-type, *clpX*, and *clpX, ftsA_G325V_*) were grown in the presence of fluorescent D-amino acids (FDAAs) which are incorporated into newly synthesized PG by the penicillin-binding proteins during transpeptidation (Kuru *et al.,* 2012). A virtual time-lapse of PG synthesis was created by first staining cells for 10 min with nitrobenzofurazan-amino-D-alanine (NADA; green but illustrated in red to improve contrast), followed by 10 min labeling with hydroxycoumarin-amino-D-alanine (HADA; blue) before imaging (Fig. 4D and 4E). Cells were imaged by SR-SIM, and progression of septal PG synthesis was analyzed by randomly picking 100 cells (two biological replicates) that had initiated septum synthesis during NADA labeling (NADA-labeled septal ingrowths not exceeding 15% of the cell diameter) and scoring cells according to the localization of the HADA signal (Fig. 4D). In the SA564 wild-type, the HADA signal localized inwards to the NADA septal signal in more than 95% of cells demonstrating that septum synthesis progresses inwards (Fig. 4D, panel i). In contrast, the HADA signal co-localized with the NADA signal in early septal ingrowths in 26 ± 1% of *clpX* cells (Fig. 4D, panel ii), while 3 ± 1% of septating *clpX* did not show any HADA signal (Fig. 4D, panel iii) supporting that inward progression of septum synthesis is arrested in a subpopulation of *clpX* cells. According to the model depicted in Fig. 1, stalling of septal synthesis precedes premature splitting and cell lysis. Therefore, HADA-localization was additionally scored in 100 cells (two biological replicates) displaying premature splitting of the unclosed NADA-stained septal ingrowths, a phenotype that is only observed in in *clpX* and *clpX*, *ftsA_G325V_* cells (Fig. 4E). Interestingly, colocalization of the HADA and NADA signals was significantly increased in *clpX* cells displaying premature splitting (P < 0.005; Fig. 4E, panel I), supporting that stalling of septum synthesis puts cells at risk for initiating daughter cell split too early. In prematurely splitting *clpX* cells, the HADA signal was also present in the lateral cell wall, showing that PG synthesis is shifted from the septal to the lateral site (Fig. 4E, panel II). Notably, stalling of septum synthesis was significantly decreased by the suppressor mutation in *ftsA* (Fig. 4D and 4E) lending further support to the idea that FtsA_G325V_ prevents lysis and restores growth of *S. aureus clpX* cells by rescuing septum synthesis.

### FtsA_G325V_ has reduced protein interactions and restores FtsA dynamics in cells lacking ClpX

In *E. coli*, FtsA interacts with early and late divisome proteins to coordinate assembly and activation of the divisome (Du and Lutkenhaus, 2017; Pichoff *et al.,* 2018). Similarly, *S. aureus* FtsA was shown to interact with multiple proteins of the divisome in a bacterial adenylate cyclase two-hybrid (BACTH) system (Steele *et al.,* 2011). To understand how the G325V amino acid substitution changed the function of FtsA, we used the BATCH BACTH system to test for interaction between FtsA_G325V_ and cell-division proteins that previously tested positive for interaction with wild-type FtsA in the BACTH system (Steele *et al.,* 2011). Using this approach, we were able to confirm all interactions of FtsA_WT_ (Fig. 5A). Interestingly, the results indicated that FtsA_G325V_ has diminished self-interaction, as well as reduced interactions with FtsZ, and several important late-stage cell division proteins including the transpeptidases PBP1, PBP3, PBP4, and LtaS, while interaction with DivIC and DivIB was unchanged (Fig. 5A). These results indicate that the FtsA_G325V_ variant cannot, or is less prone, to polymerize and to interact with many *S. aureus* divisome proteins. Based on this result, we speculated that the FtsA_G325V_ variant may compensate for the ClpX unfoldase by destabilizing divisomal protein-protein interactions that needs assistance from ClpX for disassembly at 30°C.

**Figure 5.**
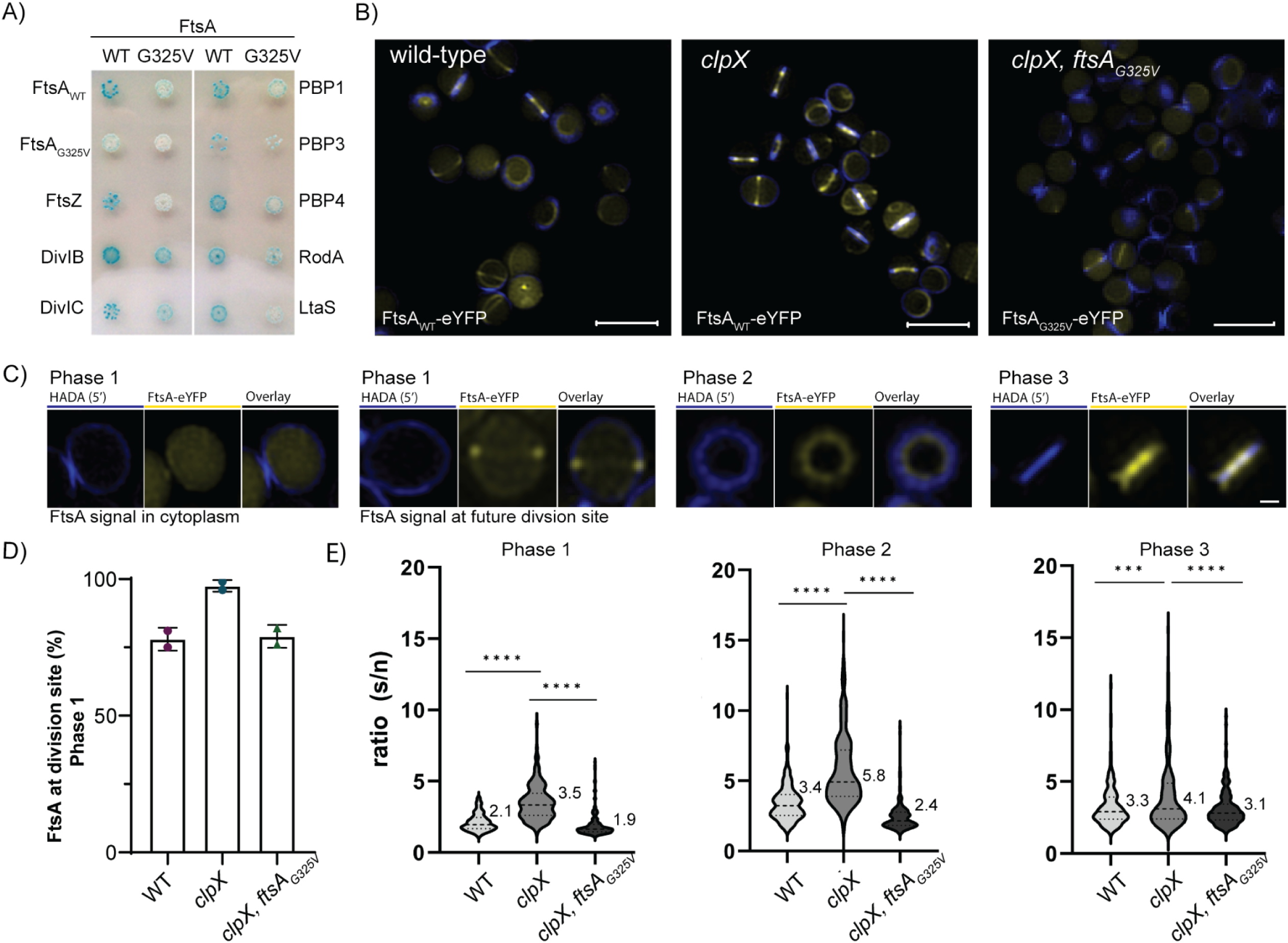
FtsA_G325V_ has reduced self-interaction and restores FtsA dynamics in cells lacking ClpX. (**A)** Interactions between FtsA_WT_ or FtsA_G325V_ (pKT25) and cell division proteins (pUT18) were assessed using BACTH: *E. coli* strain BTH101 cells were co-transformed with plasmids encoding the indicated proteins fused to domains (T25 and T18) of adenylate cyclase. Transformants were grown on selective plates containing 100 µg ml^-1^ ampicillin, 50 µg ml^-1^ kanamycin, 40 µg ml^-1^ X-Gal, and 0.5 mM IPTG. Plates were incubated at 30°C for 48 h before imaging; the image is a representative of three biological replicates. (**B**) Representative SR-SIM images showing localization of eYFP-tagged FtsA_WT_/ FtsA_G325V_ (yellow) as indicated in *S. aureus* cells that prior to imaging were stained with HADA (blue) to show localization of PG-synthesis: scale bar, 2 µM. (**C**) Example images showing localization of eYFP-FtsA signal in wild-type cells with no septal HADA signal (predivisional cells, phase 1), HADA signal in unclosed septal ingrowth (phase 2), or in closed septal ingrowths (phase 3); scale bar, 0.5 µM. (**D**) The percentage of phase 1 cells displaying the eYFP-FtsA signal at the future division site was estimated in 200 randomly picked phase 1 cells (based on the HADA signal being entirely in the lateral wall) from each of the three strains, .two biological replicates .(**E**) Septal (S) / non-septal (n) ratio of FtsA-eYFP signals were determined in 100 cells from each of the three strains in each of the three phases. Statistical significance was determined with unpaired two-tailed Student’s t tests. Statistical significance was denoted as ***P ≤ 0.001; ****P ≤ 0.0001.

To investigate the idea that ClpX becomes critical for disassembly of an FtsA-containing divisomal complex at 30°C, we next used fluorescently tagged versions of FtsA (FtsA_WT_-eYFP or FtsA_G325V_-eYFP) to visualize FtsA localization during the cell cycle in wild-type and mutant cells. Strikingly, the images revealed visual differences in the cellular eYFP signal between the three strains, with the FtsA-eYFP appearing less dispersed and more confined to the septal region in the *clpX* mutant as compared to the FtsA-eYFP signal in wild-type cells and the FtsA_G325V_-eYFP signal in the suppressor strain (Fig. 5B). To follow FtsA-localization during the cell cycle, cells were incubated with HADA for 5 min prior to imaging and the localization of FtsA was assessed in predivisional phase 1 cells (HADA signal only in peripheral wall), septating phase 2 and phase 3 cells (HADA-signal in unclosed and closed septal ingrowths, respectively) as illustrated in Fig. 5C. In phase 1 cells, two patterns of FtsA localization were observed with the FtsA signal being either evenly dispersed in the cell, or, being enriched at the future division site as illustrated in Fig. 5C and quantified in Fig. 5D. This staining pattern indicates that FtsA re-distributes to the cytoplasm before assembling at the future division site in newly born daughter cells. Of note, deletion of the *clpX* gene, significantly increased the ratio of phase 1 cells having FtsA localizing at the future site while the *ftsA s*uppressor mutation restored the distribution back to wild-type levels (Fig. 5D). To quantify the distribution of the FtsA signal between the septal and cytoplasmic site in different stages of the cell cycle, we next measured the fluorescence intensity at the septal (s) and the non-septal (n) areas and calculated the s/n ratio for cells in phase 1, 2, and 3 (Fig. 5E). Interestingly, this quantitative analysis confirmed that the ratio of FtsA localizing to the septal site was significantly increased in *clpX* cells in all three stages of the cell cycle (Fig. 5E). We conclude that ClpX directly or indirectly affects FtsA dynamics during the cell cycle and that FtsA is enriched at the septal site in cells devoid of ClpX. Interestingly, the ratio between septal and non-septal FtsA_G325V_-eYFP signal followed a similar distribution as observed for FtsA_WT_-eYFP in wild-type cells, indicating that the suppressor mutation in *ftsA* restores FtsA dynamics (Fig. 5E).

### The ftsA_G325V_ allele is conditionally lethal in a wild-type background

We next asked how the FtsA_G325V_ variant would impact growth in *S. aureus* cells having a functional ClpX. To answer this question, a wild-type *clpX* allele was back-transduced into the 564-30-8 suppressor strain by selecting for co-transduction of an *ermB* marker localized 8 kb downstream of *clpX*. Plating at different temperatures, revealed that the *ftsA_G325V_* allele conferred a heat-sensitive phenotype, as the SA564 *ftsA_G325V_* strain was unable to form colonies at 42°C while forming colonies of similar size as the wild-type cells at 30°C and 37°C (Fig. 6).

**Figure 6.**
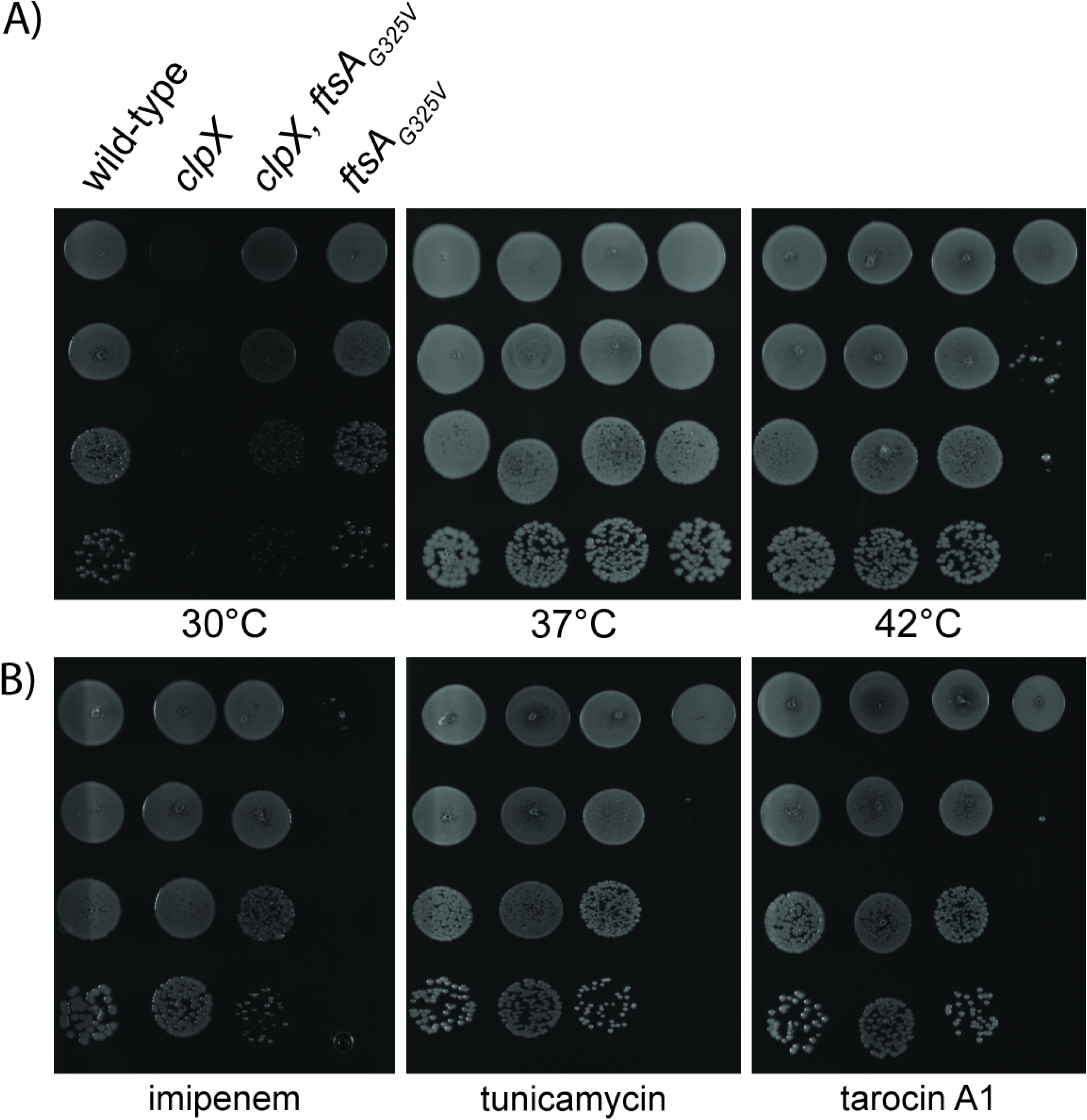
Expression of FtsA_G325V_ renders wild-type cells temperature sensitive and hypersensitive towards β-lactams and inhibitors of TarO. The *S. aureus* wild-type cells and the indicated mutants were grown exponentially in TSB at 37°C. At OD_600_ = 0.5, the cultures were diluted 10^1^ to 10^4^-fold, and 10 μl of each dilution was spotted on TSA plates supplemented with the β –lactam imipenem (0.01 µg mL^-1^), and the TarO inhibitors tunicamycin (0.5 µg mL^-1^) and tarocin A1 (1 µg mL^-1^) as indicated. The plates were incubated at 30°C, 37°C, or 42°C for 24 h.

We previously discovered that sub-lethal doses of β-lactam antibiotics rescue septum synthesis and prevent spontaneous lysis of *S. aureus clpX* mutants (Jensen *et al.,* 2019a). The only non-β-lactam compounds that stimulate the growth of the *clpX* mutant are tunicamycin and tarocin A1, two well-characterized inhibitors of the first step in the wall teichoic acid (WTA) biosynthesis (Campbell *et al.,* 2011; Lee *et al.,* 2016; Jensen *et al.,* 2019b). Strikingly, the SA564 ftsA_G325V_ strain was unable to form colonies in the presence of imipenem, tunicamycin and tarocin A1 in concentrations that rescue growth of S*. aureus clpX* mutants (Fig. 6), demonstrating that the FtsA_G325V_ variant is synthetically lethal with sub-lethal doses of β-lactams and WTA inhibitors in *S. aureus* with a functional ClpX. The synthetic lethality between the FtsA_G325V_ variant and compounds that rescue growth of the *clpX* mutant point to a functional link between FtsA_G325V_, WTA inhibitors, and β-lactam antibiotics.

### Expression of FtsA_G325V_ in wild-type cells accelerates the cell cycle and increases the risk of daughter cell lysis

To examine how the *ftsA*_G325V_ allele changes the cell morphology in a wild-type background, cells were imaged with SR-SIM (membranes stained with Nile red), TEM and scanning electron microscopy (SEM). The images revealed that the SA564 ftsA_G325V_ cells were almost twice as big as wild-type cells (1.7 ± 0.6 μm^3^ vs. 0.72 ± 0.19 μm^3^, P < 0.0001) (Fig. 7A; Supplemental Fig. S2). Furthermore, wild-type cells expressing the FtsA_G325V_ variant displayed a high frequency (53 ± 1%) of asymmetrical ingrowth of the septal cross walls (Fig. 7B; Supplemental Fig. S2, white arrows). This finding demonstrates that the *ftsA*_G325V_ allele confers this phenotype also in cells with functional ClpX. Strikingly, SEM and TEM images revealed that cultures of SA564ftsA_G325V_ contained a high number of lysed cells (Supplemental Fig. S2), and that in pairs of *ftsA_G325V_* daughter cells, one daughter cell often appeared “punctured” (Fig. 7C; Supplemental Fig. S2, white boxes). To quantify cell lysis, DNA was stained with Hoechst 33342 and PI that in contrast to Hoechst 33342 only enters cells with compromised membranes. This quantitative analysis estimated that 12 ± 3% of *ftsA_G325V_* cells were PI positive, a phenotype that was only observed in only 1 ± 0.5% of wild-type. Furthermore, the Hoechst signal appeared brighter and more compact in 10 ± 0.2% of *ftsA_G325V_* cells, indicating chromosome condensation in these cells (Fig. 7D). DNA condensation may precede lysis, as this phenotype was previously observed in *S. aureus* exposed to nisin, an antibacterial peptide that causes rapid cell lysis (Jensen *et al.,* 2020). Finally, 4 ± 0.3% of cells expressing the FtsA variant (0.25 % of wild-type cells) were neither stained with PI or Hoechst 33342, indicating that the chromosomal DNA has leaked out of the cell during cell lysis.

**Figure 7.**
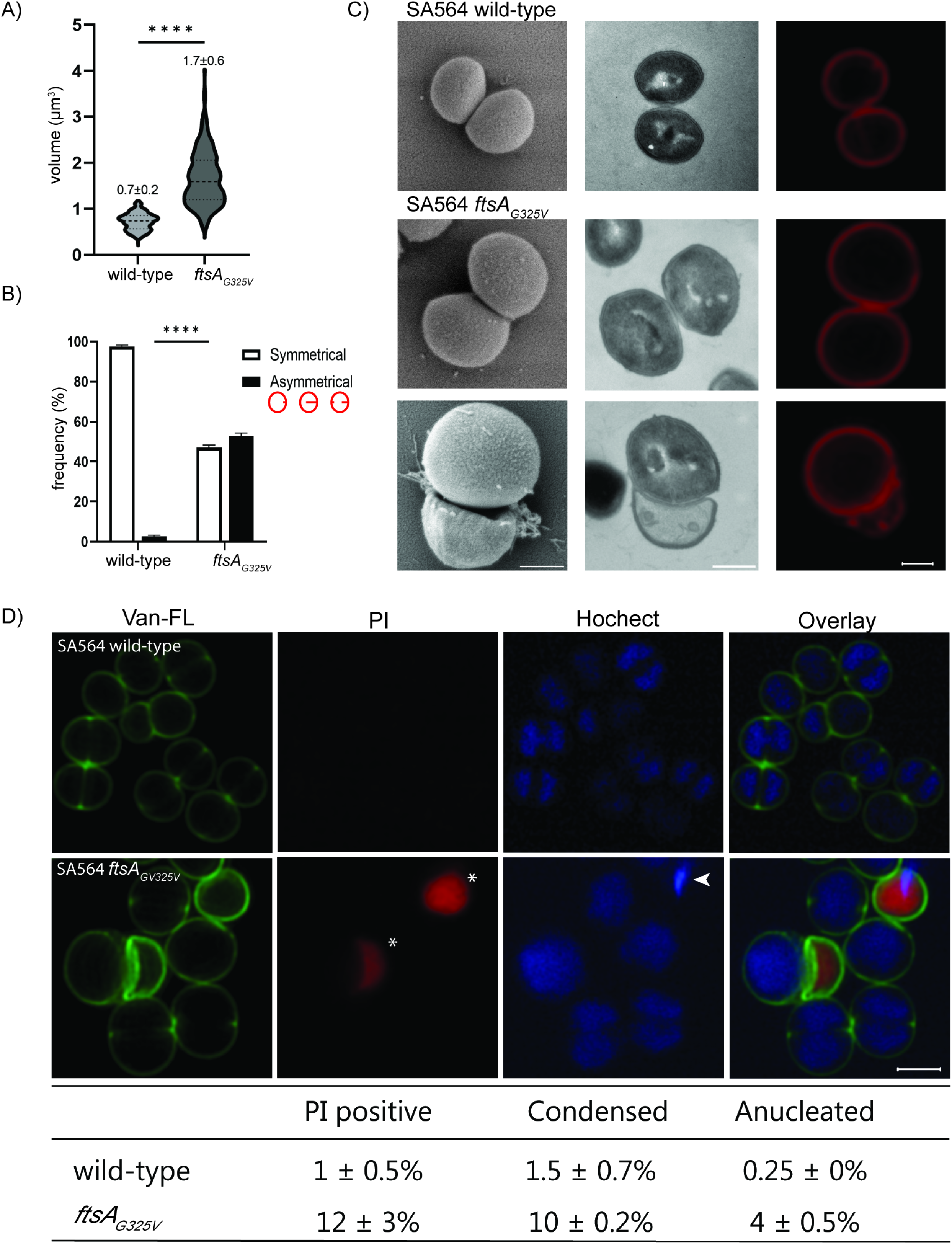
FtsA_G325V_ increases cell size, causes aberrant septum synthesis, impairs chromosome segregation and induces cell lysis. (**A+B**) To estimate the cell size and the abundance of cells displaying asymmetrical septal ingrowth cells were stained with Nile-Red prior to imaging by SR-SIM. (**A**) The volume of 300 cells representing 100 cells from each of the three growth phases were estimated by fitting an ellipse to the border limits of the membrane. (**B**) Septal ingrowths were scored as being symmetrical or asymmetrical in 100 septating cells. Both analyses were performed in three biological replicates. Statistical significance was determined with unpaired two-tailed Student’s t tests or Chi-square test of independence. Statistical significance was denoted as **** P < 0.0001. (**C**) SEM, TEM, and SR-SIM images of typical SA564 wild-type or SA564 ftsA_G325V_ cells grown to mid-exponential phase at 37°C. Scale bar, 0.2 μm. (**D**) Representative SR-SIM images of mid-exponential SA564 wild-type and SA564 ftsA_G325V_ (37°C). Prior to imaging cells were stained with PI (DNA in lyzed cells, red), Van-FL (cell wall, green), and Hoechst (DNA in all cells, blue). Cells being PI-positive (*), displaying a condensed Hoechst signal (arrowhead) or being anucleate were scored as the mean and SD from two biological replicates (lower panel). Scale bars, 2 μm.

To further investigate how the FtsA_G325V_ variant affects the cell cycle, we imaged Nile Red-stained SA564 wild-type and SA564ftsA_G325V_ cells every 5 minutes for 1 hour at room temperature (Fig. 8A). In total we followed the growth of 345 wild-type cells and 186 *ftsA_G325V_* cells. Intriguingly, the time-lapse experiment demonstrated that cells expressing the FtsA_G325V_ variant completed the cell cycle significantly faster than wild-type cells as only 2% of wild-type cells completed a full round of cell division during the 60 min experiment as compared to 72% of cells expressing the mutated FtsA variant (P < 0.0001). Examples are shown in Fig. 8A, where the four *ftsA_G325V_* cells that at T=0 were either predivisional or in an early stage of septum synthesis, at T=45 min all had completed daughter cell splitting, and at T = 60 one cell had even completed two divisions (see boxed cell in Fig. 8A). In contrast, none of the four wild-type cells that at T=0 displayed early septal ingrowth had completed the cell cycle at T=60 (example, see boxed cell in Fig. 8A).

**Figure 8.**
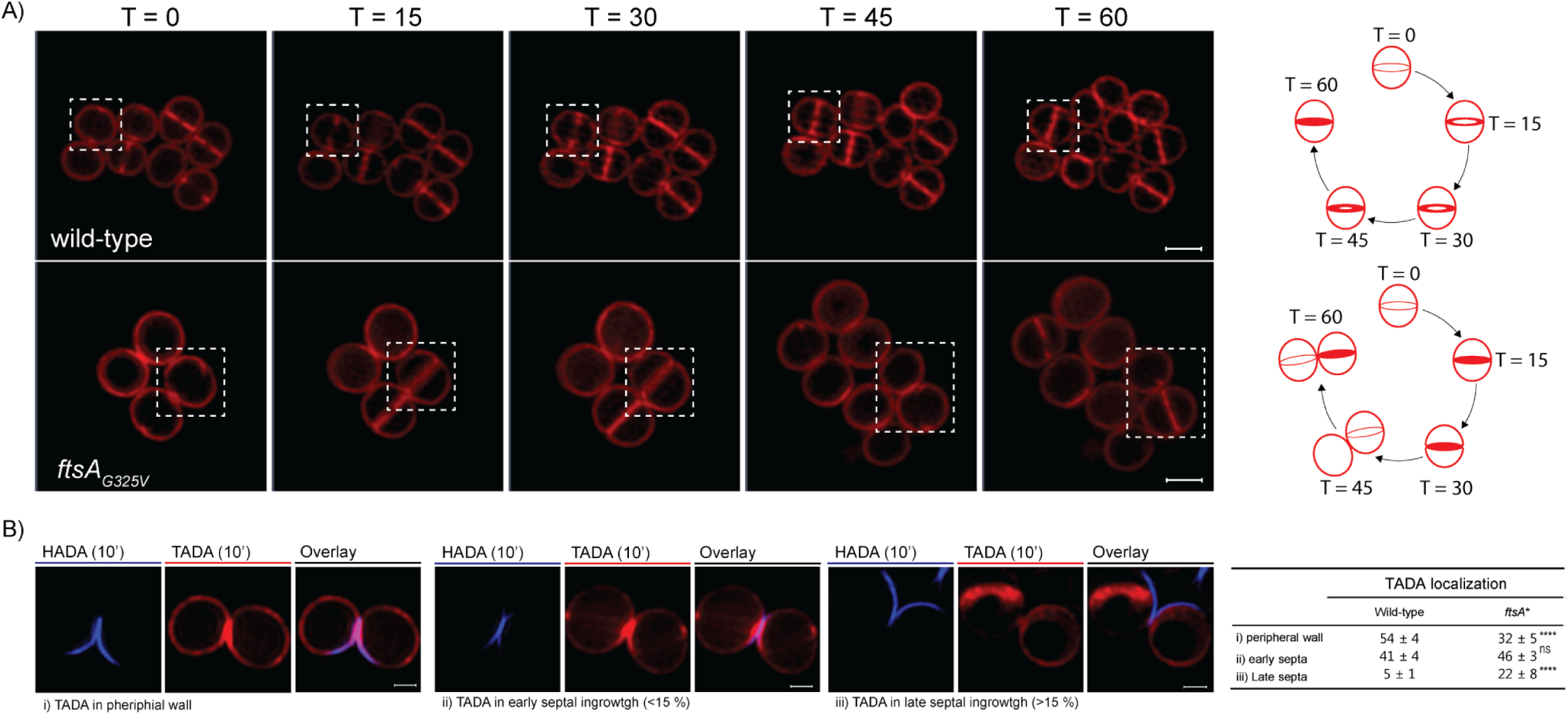
FtsA_G325V_ accelerates the cell cycle in *S. aureus.* (**A**) Still images from SR-SIM time-lapse microscopy of Nile Red stained *S. aureus* wild-type cells and cells expressing the FtsA_G325V_ variant grown at room temperature. The cell cycles of the boxed cells are depicted to the left; scale bar 1 μm. (**B**) PG synthesis was followed by incubating cells with HADA (blue) for 10 minutes followed by washing and labeling with TADA (red) for 10 min prior to SR-SIM imaging. (i-iii) Examples of TADA localization in cells that proceeded to septum closure during HADA labeling: (i): TADA signal only in peripheral wall; (ii): TADA-signal localizes at the peripheral wall and in tiny peripheral septal ingrowths (septum progresses less than 15% of the mid-cell diameter); (iii): TADA-signal localizes in septal ingrowths that have progressed more than 15% of the mid-cell diameter. The fraction of newly separated cells displaying the TADA labeling pattern depicted in i-iii was assessed in 100 cells in two biological replicates. **** P < 0.0001; ns = non-significant. Statistical analysis was performed using the Chi-square test for independence.

Above we showed that the FtsA_G325V_ antagonized the stalling of early septal ingrowth in *clpX* cells. Next, we asked how the FtsA_G325V_ affected initiation of septal PG synthesis by sequentially labeling PG-synthesis with HADA (blue) and TADA (red) and assessing localization of the TADA signal in daughter cells that had completed septum synthesis during the HADA labeling period (Fig. 8B). According to the general paradigm, newly separated *S. aureus* cells synthesize PG primarily in the peripheral wall (Monteiro *et al.,* 2015). Consistent with this paradigm, TADA was incorporated exclusively in the peripheral wall in 54 ± 4% of newly separated wild-type cells, while 46 ± 4% also displayed the TADA signal in septal ingrowths at the future division site (Fig. 8B). In comparison, the fraction of newly separated *ftsA_G325V_* cells that had initiated a new round of septum synthesis was significantly increased to 68 ± 5 % (P < 0.001), and the septal ingrowths had progressed further inwards (22 ± 8% as compared to 5 ± 1 % of wild-type cells; P < 0.001), as shown in fig. 8B. Collectively these data show that the FtsA_G325V_ accelerates the initiation of septum synthesis and that the shortened cell cycle is associated with an increased risk of daughter cell lysis.

### Suppressor mutations rescuing growth of SA564 ftsA_G325V_ activates ClpX

We next exploited the high-temperature sensitivity of the *ftsA_G325V_* mutant to select for suppressor mutations that could provide more information about the molecular mechanisms underlying this phenotype. Suppressor mutants were selected by plating SA564ftsA_G325V_ at the non-permissive temperature (42°C) for 24 hours. Large colonies containing potential suppressor mutations were independently isolated and re-streaked to confirm growth at 42°C. Four of these suppressors were selected for genome sequencing. Interestingly, two of the suppressors harbored SNPs that seemingly inactivate ClpX: in suppressor 1, ClpX expression was abolished by insertion of a single nucleotide at position 785 of *clpX* resulted in a premature stop codon at position 271 (Fig. 9A). In contrast, suppressor 2 expresses elevated level of ClpX (Fig. 9A), however, the protein is predicted to be non-functional as the mutation introduces a G150V substitution of the first G residue in the crucial GYVG loop (or pore-1 loop) involved in binding and unfolding of substrates that is conserved in ClpX proteins across all kingdoms, Fig. 9B (Martin *et al.,* 2008a & b; Siddiqui *et al.,* 2004; Rodriguez-Aliaga *et al.,* 2016). Suppressors 3 and 4 had wild-type *clpX* but deviated from the parental SA564ftsA_G325V_ strain by, respectively, a synonymous SNP at position 358 in the *uvrB* gene encoding excinuclease ABC subunit B, and a SNP in the *xdrA* gene (SA1665) resulting in an R62I substitution in a transcriptional regulator that controls genes involved in virulence and cell wall homeostasis (table S4). The selection of mutations that inactivate ClpX lends further support to a functional link between FtsA and ClpX in coordinating a critical, temperature-dependent step in *S. aureus* cell division.

**Figure 9.**
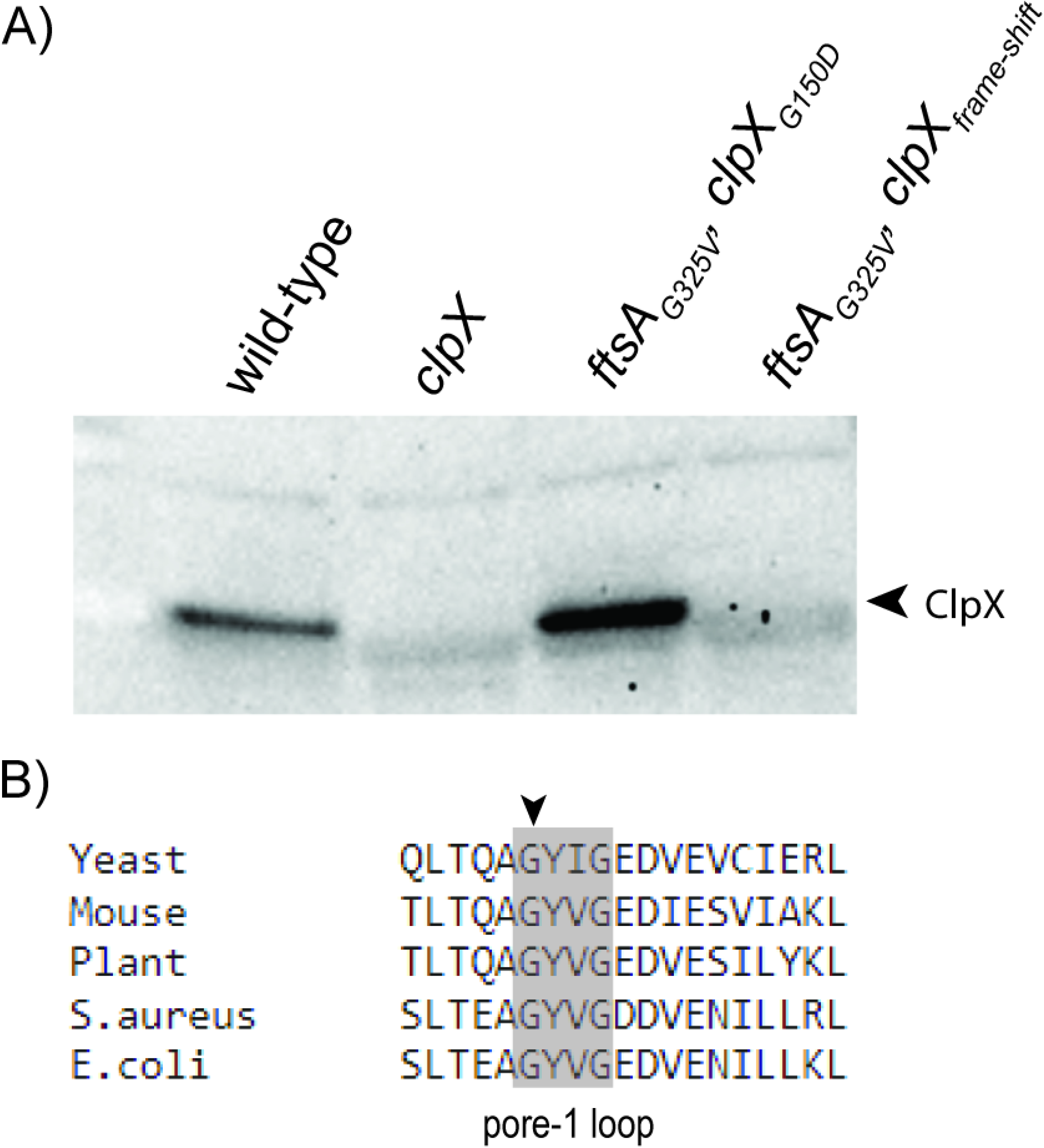
FtsA_G325V_ is selective for mutations that inactivate ClpX. (**A**) ClpX levels in cellular extracts were determined in exponential cultures of SA564 wild-type, *clpX, ftsA_G325V_* suppressor 2 (*clpX*_G150D_), and *ftsA_G325V_* suppressor 1 (*clpX*_frame shift_) by western blotting in two biological replicates. (**B**) Amino acid sequence alignment of representatives of ClpX from different organism; yeast, mouse, plant, *S. aureus,* and *E. coli.* The highly conserved pore-1 loop are highlighted in grey, and G150 is marked with an arrowhead.

## Discussion

The ClpX unfoldase localizes at the septal site (Jensen *et al.,* 2019b) and we here provide genetic evidence that the ClpX unfoldase is functionally linked with bacterial septum synthesis by showing that a G325V amino-acid substitution in the essential FtsA cell division protein rescued septum synthesis and restored the ability of *S. aureus clpX* mutants to grow at 30°C. The substituted G325 and the neighboring G324 are conserved across members of the entire actin protein family and are predicted to localize in the hydrophobic binding pocket for ATP (Fig. 2C and 2D). Point mutations in G325 (corresponding to FtsA G337 in *E. coli*) has, to the best of our knowledge, not been described before, however, a glycine to aspartate substitution in the neighboring glycine residue (G336) abolished ATP binding and conferred a complete loss of FtsA function in *E. coli* (Sánchez *et al.,* 1994; Herricks *et al.,* 2014). From the structure of FtsA, the substitution of G325 with the larger valine residue is predicted to interfere with ATP binding of *S. aureus* FtsA (Fig. 2D). The specific selection of the same G325V substitutions in two different *S. aureus* strain backgrounds, rather than selection of unspecific or loss-of-function mutations, suggest that FtsA_G325V_ must retain some functionality to rescue the growth of *S. aureus clpX* mutants. Consistent with this assumption, the mutant *ftsA* allele was tolerated in *S. aureus* cells with a functional ClpX when growing at 37°C and below. In *S. aureus* wild-type cells, introduction of the *ftsA_G325V_*allele, however, resulted in an accelerated cell cycle, asymmetrical septal ingrowths, and daughter cell lysis. Further, the FtsA_G325V_ variant was lethal in *S. aureus* growing at high temperatures. Similarly, thermosensitive alleles of the *ftsA* gene in *E. coli* map to residues in, or adjacent, to the ATP binding site (Sánchez *et al.,* 1994; Herricks *et al.,* 2014; Busiek and Margolin, 2015). In *S. aureus*, the lethality associated with the *ftsA_G325V_* allele at 42°C was selective for spontaneous suppressor mutations inactivating ClpX. The mutual, temperature-dependent suppression of lethality observed for FtsA_G325V_ and inactivated ClpX demonstrate that FtsA and ClpX are functionally linked to a critical, temperature-sensitive step in *S. aureus* cell division. *S. aureus clpX* cells can initiate septal PG synthesis, but tend to stall at an early stage with PG synthesis being redirected to the lateral wall (Fig. 4), we speculate that ClpX unfoldase activity is needed to stabilize an activated divisomal complex at 30°C, but not at 37°C. Based on the specific selection of the FtsA_G325V_ variant with reduced protein interactions, and in keeping with a conserved role of AAA+-type unfoldases in disassembly of multi-subunit complexes (van den Boom and Meyer, 2018), we propose that ClpX stabilizes the activated divisome by antagonizing FtsA protein interactions. In the most straight-forward scenario, spontaneous disassembly of FtsA polymers can be envisioned to occur at a sufficiently high pace at elevated temperatures, whereas disassembly requires assistance from the ClpX unfoldase when the temperature is lowered and protein-protein interactions become less dynamic (Li and Moore 2020). Importantly, conversion of FtsA polymers to a monomeric state was suggested to play a crucial role in the hierarchical recruitment of late division proteins and in activation of the septal PG synthesis machinery in *E. coli* (Pichoff *et al.,* 2012; Pichoff *et al.,* 2015; Du *et al.,* 2016).

Furthermore, FtsA variants deficient in self-interaction are thought to mimic an activated state of FtsA because they can bypass the need for multiple essential components of the divisome and seem capable of directly activating the septal FtsW glycosyltransferase in *E. coli* (Busiek and Margolin, 2015; Park *et al.,* 2021). Such activated FtsA variants have mutations in residues in the FtsA-FtsA interface rather than in the ATP-binding domain and, therefore, most likely change FtsA functions differently from the FtsA_G325V_ variant described here. Nonetheless, we show here that the G325V substitution in FtsA dramatically accelerated the *S. aureus* cell cycle and seemingly bypasses a key checkpoint that coordinates septum progression with activation of septal autolysins.

The increased off-septal localization of FtsA_G325V_ could be a consequence of the reduced interaction with FtsZ. Alternatively, FtsA_G325V_ may more directly interfere with attachment to the septal membrane as ATP binding was shown to induce attachment of the C-terminal amphipathic helix of FtsA to the membrane in *Streptococcus pneumonia* (Krupka *et al.,* 2014). Collectively, our data in combination with canonical models of bacterial cell division led us to propose the working model depicted in Fig. 10. According to this model, disassembly of FtsA is required for PG-synthesis to proceed to septum closure in *S. aureus*. At 30°C, ClpX unfoldase activity becomes critical for disassembly of FtsA polymers explaining why *clpX* mutants can complete septum synthesis at 37°C but not at the lower temperature. The requirement for ClpX can, however, be bypassed by the FtsA_G325V_ variant with reduced self-interaction that may be functionally related to *E. coli* activated FtsA variants. When expressed in wild-type cells, the activated FtsA_G325V_ bypasses an early cell division check-point that in wild-type cells ensures symmetrically ingrowths of septal discs, and coordinate septum synthesis with activation of septal autolysins.

**Figure 10.**
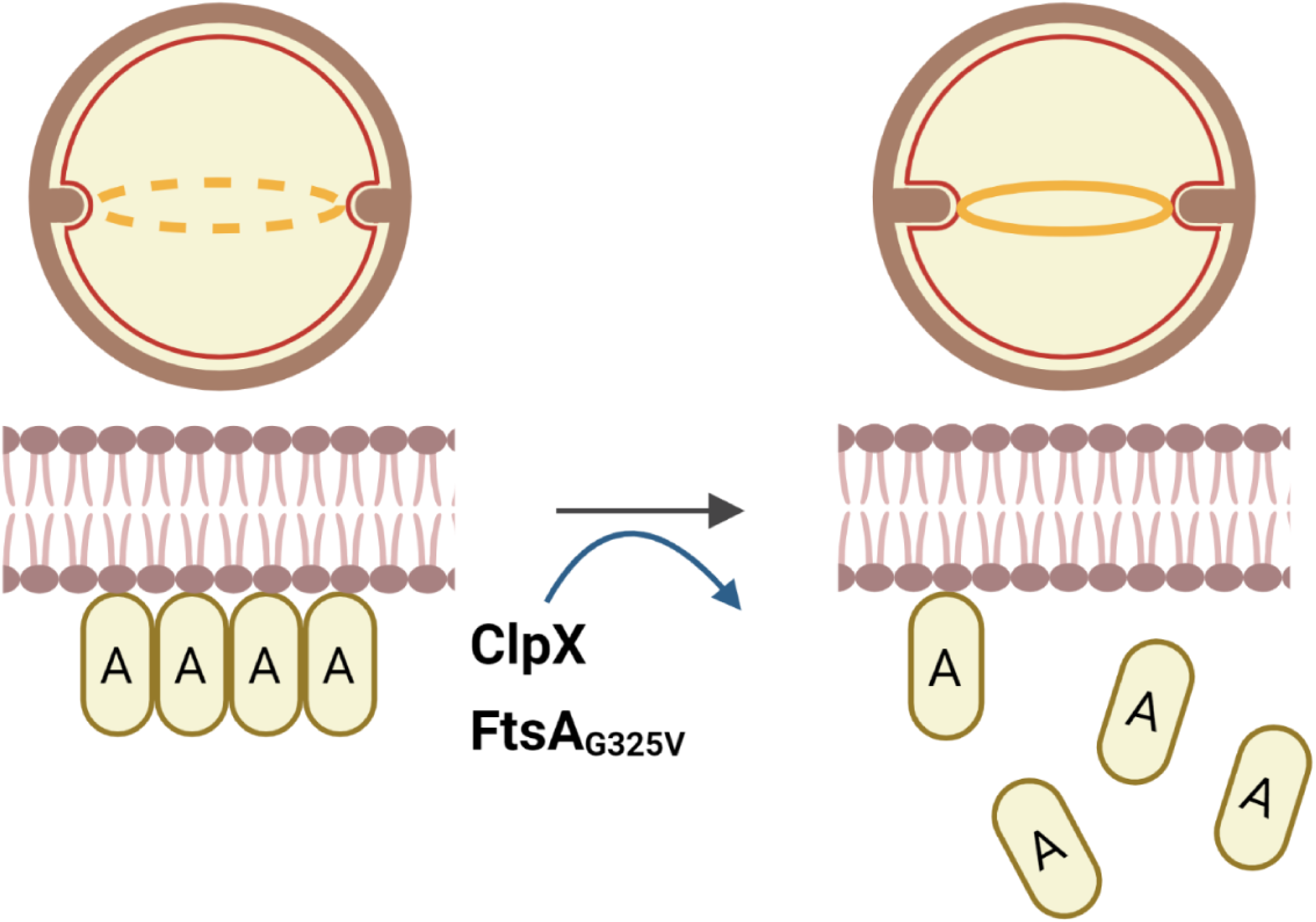
Model: ClpX promotes bacterial cell division by stimulating FtsA disassembly. Disassembly of FtsA polymers is thought to control initiation of bacterial septum synthesis (Pichoff *et al.,* 2012; Pichoff *et al.,* 2015; Du *et al.,* 2016). We propose that disassembly of FtsA polymers occurs spontaneously at 37°C, while disassembly of FtsA polymers requires assistance from the ClpX unfoldase when the temperature decreases and protein rearrangements become less dynamic. Accordingly, the redistribution of FtsA from the septal site to the cytoplasm is abrogated in cells devoid of ClpX activity. The requirement for ClpX can, however, be bypassed by the FtsA_G325V_ variant with reduced self-interaction that may be functionally related to *E. coli* activated FtsA variants as expression of the FtsA_G325V_ in ClpX-positive cells accelerates initiation of septum synthesis. See text for details.

Despite that cell division requires strict spatiotemporal control of assembly, remodeling, and disassembly of the divisome complex, the contribution of molecular chaperones to this essential process remains largely unexplored. The results presented here suggest that the ClpX unfoldase has a critical role in controlling stability of the active divisome by antagonizing FtsA protein interactions in *S. aureus*. ClpX is essential, or conditionally essential, in diverse species such as *Streptomyces*, *Caulobacter, Synechococcus, Streptococcus, and Lactococcus* (Clarke *et al.,* 1998; Jenal & Fuchs, 1998; Frees *et al.,* 2007, Piotrowski *et al.,* 2009). In *Caulobacter crescentus*, lethality of the *clpX* mutation is associated with a critical role for ClpXP in cell cycle regulation (Jenal & Fuchs, 1998). In the rod-shaped bacterial model organisms of cell division, *E. coli* and *B. subtilis*, ClpX is not essential, and inactivation of *clpX* is not reported to confer cell division defects. Nonetheless, ClpX was shown to negatively affect FtsZ polymerization *in vitro* and Z-ring formation *in vivo* in *B. subtilis*, *E. coli* and this was also observed in *Mycobacteria tuberculosis* (Weart *et al.,* 2005; Haeusser *et al.,* 2009; Dziedzic *et al.,* 2010; Sugimoto *et al.,* 2010). Therefore, the role of ClpX in controlling stability of the divisome could be widely conserved across bacterial species.

## Material and Methods

### Bacterial strains and growth conditions

Strains and plasmids used in this study are listed in S1 Table. *S. aureus* strains were cultured in tryptic soy broth (TSB) at the indicated temperatures with aeration at 180 rpm, or on TSB medium solidified with 1.5% (w/v) agar. *E. coli* strains were cultured in Luria broth (LB) at the indicated temperatures, or on LB medium solidified with 1.5% (w/v) agar. When inoculating *clpX* deletion strains, care was taken to avoid visibly larger colonies containing potential suppressor mutants.

### Construction of strains and plasmids

Primers used in this study are listed in S2 Table. KB1250 was constructed by inserting an *ermB* marker into strain 8325-4 eight kb from *clpX* between the convergently transcribed genes with locus tags SAOUHSC_1768 and SAOUHSC_1769. KB1250 was constructed as detailed in Bæk *et al*. (2014) for KB1239, but with *ermB* transcribed in the opposite direction.

The *ftsA_G325V_* single mutants were constructed by transducing the wild-type *clpX* allele (*ermB*) from KB1250 into SA564clpX, ftsA_G325V_ or 8325-4clpX, ftsA_G325V_ using phage 85 and selecting on TSA plates containing 5 μg ml^-1^ erythromycin. Replacement of *ΔclpX* with the wild-type *clpX* allele was confirmed by PCR using the ClpX_F and ClpX_R primer pair. To confirm that the complemented strains did not contain any additional mutations, PCR confirmed single mutants were sequenced on an Illumina Hi-Seq 2000 instrument at Statens Serum Institut (SSI, Denmark). Sequencing reads were analyzed using Geneious Prime @2020.2.4. The sequence analysis confirmed that SA564ftsA_G325V_ deviated from SA564 wild-type by a single mutation in *ftsA* resulting in the G325V substitution in FtsA and carried an *erm* marker eight kb from *clpX*. Additional mutations mapped to the chromosome of 8325-4 suggesting that the additional SNPs originate from KB1250 that was used as the donor in the transduction. Sequence reads of 8325-4ftsA_G325V_ revealed several additional mutations, eliminating the strain for further analysis. The SA564 (NZ_CP010890) and 8325-4 (NC_007795) genomes were used as the reference genomes.

For bacterial adenylate cyclase two hybrid (BACTH) interaction studies, *ftsA* and *ftsA_G325V_* were amplified from chromosomal DNA purified from the SA564 wild-type strain and the suppressor strain SA564clpX, ftsA_G325V_, respectively, using the primers GLUSh302F3’ and GLUSh302F5’ (Steele *et al.,* 2011). The PCR products and pUT18C were digested with KpnI and PstI and ligated, yielding plasmids pKB1382 (T18-*ftsA* fusion) and pKB1384 (T18-*ftsA_G325V_* fusion). The plasmids were maintained in *E. coli* XL1-Blue.

pCQ11:FtsA-eYFP and pCQ11:FtsA_G325V_-eYFP plasmids were constructed by amplifying FtsA and FtsA_G325V_ from chromosomal DNA isolated from SA564 wild-type and SA564clpX,ftsA_G325V_ respectively, using pCQ11-ClpX_F & pCQ11-ClpX_R (S2 table). The PCR fragment and the pCQ11 plasmid (Liew *et al.,* 2011) was digested with NheI and NcoI and ligated. The plasmids were maintained in the *E. coli* IM08B strain. To create strains containing the protein fusion pCQ11:FtsA-eYFP and pCQ11:FtsA_G325V_-eYFP were transformed into relevant *S. aureus* strains as previously described (Jensen *et al.,* 2019a).

### Genetic suppressor analysis and SNP detection

Genetic suppressor analysis was performed as previously described (Bæk *et al.,* 2016). Small colonies of SA564 and 8325-4 *clpX* mutants were spread on TSB plates and incubated at 30°C or 37°C. Colonies that appeared visibly larger than the background colonies were purified by restreaking twice at the appropriate temperature. One of the obtained suppressor mutants in the SA564 *clpX* genetic background (564-30-8) was chosen for further analysis and genomic DNA was prepared for WGS. WGS was performed on an Illumina Hi-Seq 2000 instrument (Fasteris, SA, Geneva, Switzerland). The sequencing reads obtained by Illumina sequencing of SA564clpX and the suppressor mutant 564-30-8 were aligned to the assembled SA564 genome (GenBank accession numbers CP010890 and CP010891 for chromosome and plasmid, respectively) using Bowtie2, version 2.2.3, default parameters (Langmead & Salzberg, 2012). The sequences have been submitted to the European Nucleotide Archive under accession number PRJEB8687 (http://www.ebi.ac.uk/ena/data/view/PRJEB8687). Alignments were converted to sorted, indexed binary alignment map (BAM) files, and SNPs and indels were called with SAMtools and BCFtools, version 1.1, default parameters (Li *et al.,* 2009). BAM files were converted to browser extensible data (BED) format, and sequencing depth was computed with BEDtools, version 2.21.0. Average read coverage was calculated to be 362 and 1585 for the chromosome and plasmid, respectively. The sequencing reads from SA564clpX or 564-30-8 covered the whole reference genome except for the expected *clpX* deletion in both strains. The SA564 reference genome was annotated with the RAST server (Aziz *et al.,* 2008), and annotations and calculated amino acid changes were assigned to the variant set using a custom R script. Variants were excluded if the quality score for the base call was < 100, or if the phred-scaled *p*-value using Fisher’s exact test for strand bias was > 200 for indels or > 60 for SNPs. The remaining variants were verified by capillary sequencing using appropriate PCR primers (S2 Table). By this method, one SNP and one indel (both in the tRNA-Leu-CAA gene) were identified between the reference genome and our SA564 wild-type, no nucleotide variations were identified between the SA564 wild-type and the SA564 *clpX* mutant, and one SNP (*ftsA*_G325V_) was identified between 564-30-8 and the SA564 *clpX* mutant.

Genomic DNA was isolated from 20 suppressor mutants selected in the SA564 and 8325-4 backgrounds. From these, the *ftsA* gene was amplified by PCR with the primer pair KB80F and KB80R and sequenced by capillary sequencing. One SNP (resulting in a G325V substitution in FtsA) was identified in *ftsA* from one of the 8325-4 *clpX* suppressor mutants (strain 83-30-Y).

To illuminate the role of the FtsA_G325V_ in *S. aureus* cell division, SA564ftsA_G325V_ cells were grown at 42°C to select for spontaneous suppressors. In short, SA564ftsA_G325V_ was grown exponentially in TSB at 37°C until OD_600_ of 0.5 ± 0.1. The cultures were diluted 10^1^ - 10^4^-fold in 0.9% (w/v) NaCl. 100 μl of each dilution was plated on TSA at 42°C for 24 h. Colonies that appeared visibly larger than the background colonies were picked and restreaked on TSA to confirm growth at 42°C. Genomic DNA was prepared from overnight cultures of four SA564ftsA_G325V_ suppressors grown at 42°C and sequenced on an Illumina Hi-Seq 2000 instrument at Statens Serum Institut (SSI, Denmark). Sequencing reads were analyzed using Geneious Prime @2020.2.4. The assembled sequences from the suppressor mutants were compared with *S. aureus* SA564ftsA_G325V_ parental strain to identify single nucleotide polymorphisms, deletions, and insertions unique to the suppressor mutants. The SA564 (NZ_CP010890) genome was used as the reference genome.

### Amino acid sequence alignment of the actin family and ClpX

To demonstrate that the G325 and the neighboring G324 are highly conserved across the entire actin protein family, FtsA (WP_000391031.1), HSP70 (WP_050960908.1), and ParM (WP_000073389.1) from *S. aureus,* MreB from *E. coli* (AAN82446.1), and human actin (NP_001092.1), were chosen for protein alignment (Clustal Omega).

To demonstrate that ClpX_G150_ and the pore-1 loop is conserved in ClpX are highly conserved ClpX from yeast (CAI5030975.1), mouse (XP_021061903.1), plant (NP_568792.1), *S. aureus* (P63790.1) and *E. coli* (HBD5768364.1) were chosen for protein alignment (Clustal Omega).

### Western blot analysis

*Whole cell extracts* were isolated from exponential *S. aureus* cultures by resuspending pellets in 50 mM Tris-HCL, pH = 8 containing 5 μg ml^-1^ lysostaphin normalized to an OD_600_ of 1 ml^−1^ followed by incubation at 37°C for 30 min. *Cell wall associated proteins* were isolated by resuspending pellet in 4% SDS normalized to an OD_600_ of 1 ml^−1^ and incubated at room temperature for 45 minutes. FtsA and Sle1 levels were detected by running samples on NuPAGE 10% Bis-Tris gels (Invitrogen) with morpholinepropanesulfonic acid buffer (Invitrogen). The proteins were subsequently blotted onto a polyvinylidene difluoride membrane (0.45-μm pore size; Invitrogen). To avoid Protein A signal, membranes were pre-blocked with human IgG. Proteins were detected using rabbit-raised antibodies against staphylococcal Sle1 (Kajimura *et al.,* 2005), ClpX (Jelsbak *et al*., 2010) or rabbit-raised antibodies against streptococcus FtsA kindly provided by Miguel Vicente’s lab. All of the Western blots were repeated at least three times with similar results. Bound antibody was detected using WesternBreeze chemiluminescent anti-rabbit kit. Densitometry analysis was performed using the ImageJ gel analysis tool.

### Zymography

Cell wall associated protein were isolated as described above. Proteins were separated by SDS-PAGE on a 10% resolving gel containing 1% heat-killed SA564 wild-type cells. The gel was washed three times in ionized water and incubated for 24 h in renaturing buffer (50 mM Tris-HCl [pH 7.5], 0.1% Triton X-100, 10 mM CaCl2, 10 mM MgCl2) at 37°C with gentle agitation. The gel was stained in a solution containing 0.4% methylene blue, 0.01% KOH, and 22% ethanol for 1 min. Excess dye was removed (1 h wash in water) prior to imaging. Experiment was performed in duplicate.

### SR-SIM

#### Image acquisition

SR-SIM analysis was carried out at the Core Facility for Integrated Microscopy (CFIM), Faculty of Health and Medical Sciences, University of Copenhagen. Images were acquired using an Elyra PS.1 microscope (Zeiss) with five grid rotations using a Plan-Apochromat 63x/1.4 oil DIC M27 objective, a Pco.edge 5.5 camera. Images were reconstructed using ZEN software (black edition, 2012, version 8.1.0.484) based on a structured illumination algorithm, using synthetic, channel specific optical transfer functions and noise filter settings ranging from −6 to −8. Laser specifications can be seen in S3 Table.

#### Live and dead staining

Exponential cultures of *S. aureus* grown at the indicated temperatures were stained for 5 min at room temperature using the DNA dyes Hoechst 33342 and PI, and a fluorescein conjugated variant of vancomycin (Van-FL) that labels the entire cell wall (S3 Table). To stain the cell wall, a mixture of equal amounts of vancomycin and a BODIPY FL conjugate of vancomycin (Van-FL, Molecular Probes) was used as previously described (Pinho and Errington, 2003). Stained cells were placed on an agarose pad (1.2% in PBS) in a Gene Frame (Thermo Fisher Scientific) and image by SR-SIM as described above. The frequency of dead cells was assessed among 300 cells for each strain in three biological replicates. Cells were scored as “live” if they displayed a Hoechst signal, but not a PI signal. Cells were scored as dead if they either displayed a Hoechst and a PI signal and if they did not display any of the two (anucleated cells). Statistical analysis was performed using the Chi-square test for independence in GraphPad Prism 9.5 (GraphPad Software LLC).

#### Analysis of the cell cycle

Exponential cultures of *S. aureus* grown at the indicated temperatures were stained for 5 min at room temperature using the membrane dye Nile Red and the cell wall dye WGA-488 (S3 Table). Cells were placed in a Gene Frame (Thermo Fisher Scientific) on an agarose pad (1.2% in PBS) and image by SR-SIM as described above. To analyze progression of the cell cycle, 300 random cells were scored according to the stage of septum ingrowth. Cell that had not initiated septal PG synthesis was assigned to phase 1, cells undergoing septal synthesis was assigned to phase 2, and cells displaying a complete septum was assigned to phase 3 as described by Monteiro *et al*. (2015). To further evaluate progression of the cell cycle in *S. aureus* cells, the fraction of phase 2 cells displaying asymmetrical septum ingrowth or showing signs of premature splitting (based on WGA-488 signal) was scored in 200 septating (phase 2) cells. All analyzes were performed on at least two biological replicates. Statistical differences was assessed by Chi-square test of independence in GraphPad Prism 9.5 (GraphPad Software LLC).

#### Progression of PG synthesis - 30°C

Progression of septal PG synthesis in normal septating cells and cells displaying premature split was analyzed by sequentially labeling cells with NADA and HADA at 30°C. In short, exponential cultures grown at 30°C were stained with NADA for 10 minutes at 30°C. Unbound NADA was removed by washing twice in PBS. Cells were subsequently stained 10 minutes with HADA. Excess HADA was removed by repeating the washing step. Cells were image by SR-SIM and progression of septal PG synthesis was analyzed in 50 cells as described previously (Jensen *et al.,* 2019a). In short, HADA incorporation was assessed in cells that had initiated septum formation during the initial labeling. The analyses were performed for two biological replicates. Note, the analysis was performed on the same data set as used for the analysis performed in Jensen *et al.,* 2019a. Here SA564clpX, ftsA_G325V_ was included in the imaging but not the analysis.

#### Analysis of FtsA-eYFP dynamics

To assess FtsA localization relative to active PG synthesis, SA564 wild-type pCQ11:FtsA-eYFP and SA564 *clpX* pCQ11:FtsA-eYFP were grown exponentially at 37°C. After four generations (OD_600_ ∼0.2), IPTG was added to a final concentration of 50 μM and cultures were grown for one hour prior to imaging. Cells were stained for 5 min at 37°C with HADA. Excess HADA was removed by washing twice in PBS. Cells were imaged by SR-SIM. Septal:non-septal ratios were determined in 100 cells from each of the three phases by drawing an intensity line through the cell. Image analysis was performed using Fiji. The analysis was perform in three biological replicates. Student’s t-test was perform in GraphPad Prism 9.5 (GraphPad Software LLC) to determine differences in fluorescence ratio.

#### Estimation of cell size

Volume was used as a measure for cells size of *S. aureus* cells based on the assumption that cell shapes is that of a prolate spheroid. Volumes were estimated as described previously (Monteiro *et al.,* 2015). For each strain, the volume of 100 cells representing each of the three growth phases were estimated by fitting an ellipse to the border limits of the membrane. Measurements of the minor and major axis were used to estimate the volume based on the equation V = 4/3πab2 where a and b correspond to the major and minor axes, respectively. Measurement were obtained using Fiji.

#### Time-lapse

To examine the effect of the FtsA_G325V_ variant on growth, SA564 wild-type and *ftsA*_G325V_ cells were analyzed by time-lapse microscopy. Prior to imaging, exponential culture grown at 37°C were stained with Nile Red. To follow growth, images were acquired every 5 minutes for 60 minutes (13 frames). Growth was followed in 345 wild-type cells and 186 *ftsA_G325V_* cells. It was determined how long (number of frames) each cell spend in the different growth phases (phase 1, phase 2, and phase 3). If a cell completed all three phases during the duration of the experiment it was considered a full cell cycle. Statistical differences was assessed by Chi-square test of independence in GraphPad Prism 9.5 (GraphPad Software LLC).

#### Progression of PG synthesis - 37°C

To further assess accelerated cell cycle in cells expressing the *ftsA_G325V_*allele, progression of septal PG synthesis was analyzed as described above with the following changes; cells were grown and strained at 37°C, cells were initially stained with HADA followed by staining with TADA (red). Cells were image by SR-SIM and progression of PG synthesis was analyzed in 50 newly separated daughter cells for each strain that had completed septum synthesis during the HADA labeling period by evaluating TADA incorporation. TADA was scored as phase 1 (TADA exclusively in the peripheral wall), early phase 2 (thin septal ring, < 15% septum ingrowth) or late phase 2 (> 15% septum ingrowth). Statistical differences was assessed by Chi-square test of independence in GraphPad Prism 9.5 (GraphPad Software LLC).

### Time lapse phase contrast microscopy

Time lapse phase contrast microscopy was performed as described previously (Jensen *et al.,* 2019a). Shortly, *S. aureus* cultures were grown exponentially at 37°C (OD_600_ = 0.1) before shifting cultures to 30°C. Washed cells were placed on TSB-polyacrylamide (10%) slides. Time-lapse microscopy was performed using a DV Elite microscope (GE healthcare) with a sCMOS (PCO) camera with and a 100x oil-immersion objective. Images were acquired with 200 ms exposure time every 6 minutes for at least 6 h at 30°C using Softworx (Applied Precision) software. Experiment was performed in triplicate.

### Transmission electron microscopy (TEM)

Exponential cells of *S. aureus* grown at 30°C were collected from a 10 mL aliquot by centrifugation at 8,000 x g, and the cell pellets were suspended in fixation solution (2.5% glutaraldehyde in 0.1 M cacodylate buffer [pH 7.4]) and incubated overnight at 4°C. The fixed cells were treated with 2% osmium tetroxide. Hereafter cells were treated with 0.25% uranyl acetate to enhance contrast. The pellets were dehydrated in increasing concentrations of ethanol, followed by pure propylene oxide. Dehydrated cells were embedded in Epon resin and thin sections were stained with lead citrate for electron microscopy. Cells were observed in a Philips CM100 BioTWIN transmission electron microscope fitted with an Olympus Veleta camera with a resolution of 2,048 by 2,048 pixels. Sample processing and microscopy were performed at CFIM.

### Scanning electron microscopy (SEM)

Exponential cultures were prepared for SEM as described previously (Jensen *et al.,* 2019a). In short pellets were fixed in 2% glutaraldehyde in 0.05 M sodium phosphate buffer, pH 7.4. Cells were sedimented on coverslips at 4°C for 1 week. Cells were washed in in 0.15 M sodium phosphate buffer, pH 7.4. Specimens were post fixed in 1% OsO_4_ (in 0.12 M sodium cacodylate buffer, pH 7.4) for two hours. Washed specimens were progressively dehydrated to 100% ethanol and critical point dried (Balzers CPD 030) with CO_2_. Cells were mounted on stubs using colloidal silver as an adhesive and sputter coated with 6 nm gold (Leica Coater ACE 200). Cells were imaged using FEI Quanta 3D scanning electron microscope operated at an accelerating voltage of 2 kV. Sample preparation and SEM imaging was performed at CFIM.

### Bacterial adenylate cyclase two-hybrid (BACTH)

pKT25/pKNT25 and pUT18/pUT18C vectors containing the genes of interest were introduced into chemically competent BTH101 cells and transformation reactions spotted onto LB agar supplemented with 100 µg ml^−1^ Amp, 50 µg ml^−^ Kan, 40 µg ml^−^ X-gal and 0.5 mM IPTG. Plates were incubated at 30°C for 48 h before images were taken and coloration assessed. Experiments were performed in triplicate and a representative result is shown.

## Acknowledgements

We would like to thank Ewa Kuninska and Vi Phuong Thi Nguyen (University of Copenhagen) for technical assistance, Sylvain Lemeille and Patrick Linder (University of Geneva) for providing the sequence for the assembled SA564 genome, the bioinformatic staff at Fasteris S.A. (Geneva) for assistance with Illumina sequencing analysis, and the staff at the Center for Integrated Microscopy (University of Copenhagen) for assistance with electron microscopy. We would also like to thank Miguel Vicente’s lab for providing the FtsA antibody. This work was in part funded by The Danish Research Council (DFF, FTP), grants 0136-00200B and 4184-00033 to Dorte Frees and the Wellcome Trust (212197/Z/19/Z to Simon J. Foster).

## Supporting information

**Figure S1. Analysis of single cells reveals that the FtsA_G325V_ variant prevents growth arrest and spontaneous lysis of *S. aureus clpX* mutants**

Images from automated phase contrast time-lapse microscopy of SA564 wild-type, *clpX*, and *clpX*, *ftsA_G325V_* cells growing at 30°C. Displayed microcolonies are representative of the typical growth among at least 20 imaged microcolonies of SA564 wild-type and *clpX, ftsA_G325V_* cells whereas the microcolony displayed for *clpX* belongs to the minority of *clpX* cells that were able to initiate growth and form a micro colony. Scale bar, 5 μm.

**Figure S2. Expression of FtsA_G325V_ in wild-type cells induces cell lysis in one daughter cell**

SR-SIM (Nile Red), SEM, and TEM overview images of SA564 wild-type or *ftsA* cells grown to mid-exponential phase at 37°C. Asterisk mark examples of cells displaying asymmetrical septal ingrowths (upper panel); white boxes in the left panel enlarge cells displaying the characteristic morphology of *ftsA*_G325V_ lysis (punctured cells) in the right panel. Scale bar, 5 μm.

